# Conformational Stability of Peroxidase from latex of *Artocarpus lakoocha*: Influence of pH, Chaotropes, and Temperature

**DOI:** 10.1101/2023.01.27.525866

**Authors:** Kirti Shila Sonkar, Manendra Pachauri, Amit Kumar, Medicherla V. Jagannadham

## Abstract

A novel heme-peroxidase has been extracted from the latex of the medicinal plant *Artocarpus lakoocha (A. lakoocha)*, known for its potential anti-inflammatory and wound healing properties. To study its stability, structure, and dynamics, this protein was analyzed using far-UV circular dichroism, fluorescence spectroscopy, and activity measurements. The results demonstrated the presence of three folding states: the **native state** (N) at neutral pH, **intermediate states** including molten globule (MG) at pH 2 and acid-unfolded (UA) at pH 1.5 or lower, and acid-refolded (A) at pH 0.5, along with alkaline denatured (UB) at pH 8-12 and the third **denatured state** (D) at GuHCl concentrations exceeding 5 M. Absorbance studies indicated the presence of free heme in the pH range of 1-2. The protein showed stability and structural integrity across a wide pH range (3-10), temperature (70 °C), and high concentrations of GuHCl (5 M) and urea (8 M). This study is the first to report multiple ‘partially folded intermediate states’ of *A. lakoocha* peroxidase, with varying amounts of secondary structure, stability, and compactness. These results demonstrate the high stability of *A. lakoocha* peroxidase and its potential for biotechnological and industrial applications, making it a valuable model system for further studies on its structure-function relationship.

## 1. Introduction

*Artocarpus lakoocha* (*A. lakoocha*) is a medicinal plant of the Moraceae family. It is a widely used medicinal plant of India for the treatment of many diseases like intestinal fluke *Haplorchis* taichui (Wongsawad, Wongsawad, Luangphai, & Kumchoo, 2005), taeniasis (Charoenlarp, Radomyos, & Bunnag, 1989), and cosmetics (Theorell, 1941). In an earlier report, we purified and characterized a novel peroxidase from *A. lakoocha.* The preliminary biophysical studies on *A. lakoocha* peroxidase revealed that the isolated novel enzyme from *A. lakoocha* is a stable heme-peroxidase with anti-inflammatory and wound healing properties (Sonkar et al., 2015). These broad applications provide a stimulus for detailed characterization of this peroxidase regarding folding behavior. Proteins undergo extensive conformational changes as part of their functionality; tracing these changes is important for understanding the mechanism by which proteins function.

Folding and conformational stability of proteins are primarily based on their amino acid sequence as well as cellular external variables such as temperature, pH, and chemical denaturants (Anfinsen, 1973; Masson & Lushchekina, 2022; Mohamed, El-Badry, Drees, & Fahmy, 2008). To predict the behavior of protein folding, it is important to understand the effect of these external variables on the thermodynamics of the folding process. Several other forces such as hydrogen bonding, disulfide bonding, entropic effects, hydrophobic, van der Waals, and electrostatic interactions, stabilize the folded state of the protein.

From unfolded chain to functional three dimensional state, the protein undergoes various non-native intermediate forms that are involved in various cell phenomena such as chaperone action (Díaz-Villanueva, Díaz-Molina, & García-González, 2015; Uversky, 2010), amyloid fibril formation in pathological cells (Almeida & Brito, 2020). A better understanding of protein folding mechanisms can help in treating diseases associated with protein misfolding, such as Alzheimer’s, Parkinson’s, and prion diseases (Rochet & Lansbury, 2000; Stefani & Dobson, 2003). The protein folding pathway is characterized by the accumulation of many intermediates between the native (N) and fully unfolded (U) states. Understanding these intermediate states is important to know how and when various forces come into play in directing the protein to attain three-dimensional functional structures. Studying these intermediates will help to explain the mechanism of folding of protein into the specific biologically active conformations (Benaim & Villalobo, 2002). Protein folding is a sequentially ordered process with the existence of stable ‘molten globule’ (MG) conformations between fully folded and unfolded states [3]. Molten globules serve as a common equilibrium intermediate state in the protein folding pathway, which is acquired under mild denaturing conditions. These molten globule states are believed to give information on early events in the folding process, at a stage when specific side-chain interactions are not yet formed. Molten globule was considered less compact than its native state and more compact than its unfolded state with prominent secondary structure but slight or no tertiary packing (Baldwin & Rose, 2013; Bhattacharyya & Varadarajan, 2013). The characteristics of the molten globule state are an exposed hydrophobic surface that binds a hydrophobic dye. This dye helps to yield experimental evidence concerning the structure of intermediates throughout the protein folding state (Judy & Kishore, 2019). The exact role of molten globule state is controversial among the proteins. The same protein may have varied molten globule states in the folding pathway (Bychkova, Dolgikh, Balobanov, & Finkelstein, 2022). The experimental techniques typically used to detect molten globule states are far-UV and near-UV CD spectroscopy, which detect the secondary and tertiary structure of protein. The hydrophobic dye, such as ANS (8-anilino-naphtalene-1-sulfonate) binding experiments, detects the formation of a loose hydrophobic core and estimates the extent of hydrophobic area exposed to the solvent.

With this background, the conformational stability of *A. lakoocha* peroxidase from the latex of *A. lakoocha* has been examined in a wide range of pH and temperature and in the presence of various chemical denaturants (GuHCl). The present analysis sheds light on the biophysical properties of *A. lakoocha* peroxidase to find the correlation between its structure and function and the root and logic behind its distinct physiochemical properties. The conformational stability of the native protein as well as intermediate states have been evaluated, compared, and discussed. Several intermediates with distinct spectroscopic properties are populated under various conditions.

Our initial biochemical analysis has revealed that the isolated peroxidase has a molecular mass of 53 kDa, and its molecular structure consists of four disulfide bridges, fourteen tyrosine residues, and seventeen tryptophan residues, with an extinction coefficient (ε280^1%^) of the enzyme at 16.3 (Sonkar et al., 2015). This peroxidase showed remarkable stability under various denaturing conditions. In this section, conformational studies, and other factors responsible for such high stability of peroxidase against various denaturing conditions, such as high concentrations of urea and guanidine hydrochloride and a wide range of pH and temperature, have been reported. Several equilibrium intermediate states with different stabilities have been obtained and characterized to increase the understanding of the folding aspects of the peroxidase molecule. The structural statistics about transition-state intermediates have been examined to identify the whole protein folding method of peroxidase from the latex of *A. lakoocha*. These studies establish a stable folded state of protein, which reflects its functional stability, and are aimed at providing an insight into the folding of proteins in general, which may find application in biotechnology and other fields of science.

## 2. Materials and methods

### 2.1 Materials

Peroxidase was purified from fresh latex of *Artocarpus lakoocha* plant using the method publish ealier (Sonkar et al., 2015). ANS (8-anilino-1-naphthalene sulphonate) and GuHCl were purchased from Sigma Chemicals, USA. Other reagents used were of analytical grade. The samples were centrifuged before spectroscopic measurements, and the exact concentration of the protein and pH of the buffer were determined before the measurements were taken.

### 2.2 Methods

#### 2.2.1 Absorbance spectroscopy

Spectroscopic analysis was performed using a Beckman DU-640B spectrophotometer with a constant-temperature cell holder. A protein concentration of 1 mM was used for all absorbance measurements. The concentration of the enzyme was determined spectrophotometrically using an extinction coefficient (ε2801%) of 16.3 M^-1^cm^-1^. Far-UV circular dichroism (CD) measurements were done using a JASCO J-815 spectropolarimeter (JASCO Corporation, Tokyo, Japan) equipped with a constant temperature cell holder. The instrument was calibrated using ammonium (+)-10-camphor sulfonate, and conformational changes in the secondary structure of the peroxidase were monitored in the 200-260 nm range. After subtracting appropriate blanks, mean residue ellipticity was calculated using the equation provided by Balasubramanian et al (Balasubramanian & Kumar, 1976) as

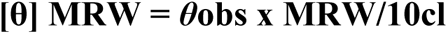

Where *θ*obs is the observed ellipticity in degrees, MRW represents mean residue weight, c represents the concentration of protein (mg/ml), and l represents the path length in centimeters. The mean weight of amino acid residues (MRW) was taken as 112. Sensitivities of 1 and 2 millidegrees/cm were used for far-UV CD measurements.

#### 2.2.2 Fluorescence spectroscopy

Fluorescence measurements were carried out using an Eclipse Cary Varian UV-Vis spectrofluorometer (Varian Inc., Palo Alto, California, USA) attached to a Peltier temperature controller. For intrinsic tryptophan fluorescence, the protein was excited at 292 nm, and the fluorescence emission was monitored in the 300-400 nm range. For ANS fluorescence, the excitation was done at 380 nm, and the emission was monitored between 400-600 nm. For all experiments, a protein concentration of 1 mM and excitation and emission slit widths of 10 and 5 nm were used, respectively. All the fluorescence spectra were recorded at 25 °C unless stated otherwise. The activity of *A. lakoocha* peroxidase under various conditions of pH and chemical denaturant was monitored using hydrogen peroxide as a hydrogen acceptor, and other compounds such as guaiacol as a hydrogen donor. Twenty micrograms of peroxidase were added to 0.2 mM of hydrogen peroxide (substrate I) with guaiacol in a 50 mM acetate buffer (substrate II). The rate of change of absorbance of guaiacol was determined at 470 nm.

#### 2.2.3 pH denaturation of A. lakoocha peroxidase

Acid denaturation of peroxidase was observed at a broad pH range using various buffers, such as KCl–HCl buffer (pH 0.5–1.5), Gly–HCl buffer (pH 2–3.5), sodium acetate buffer (pH 4.0–5.5), sodium phosphate buffer (pH 6.0–8.0), Tris–HCl buffer (pH 8.5–10.5), and Gly–NaOH buffer (pH 11–12.5). The total concentration of all the buffers used in this study was 50 mM. A stock solution of the protein was added to the appropriate buffer, and the mixture was incubated for 24 hr at 25 °C. The pH of the final suspension and the concentration of the protein in each sample were confirmed by measurement at the final steps.

#### 2.2.4 8-anilino-1-naphthalene sulfonate (ANS) binding assay

ANS is highly soluble in methanol (Hawe, Sutter, & Jiskoot, 2008), and a minimal amount of methanol does not affect the structure of the protein, as *A. lakoocha* peroxidase shows activity and stability up to 50 % methanol (Sonkar et al., 2015). Based on this, ANS solution was prepared in methanol. An extinction coefficient value of 5000M−1 cm−1 at 350 nm was used for ANS concentration measurement. A molar ratio of 1:100 for protein to ANS was used, and the suspension was incubated for 30 minutes in the dark. Subsequently, the extrinsic fluorescence of the protein-ANS complex was monitored by recording the emission spectrum between 400-600 nm using an excitation wavelength of 380 nm. Protein concentration of 1 mM and slit widths of excitation and emission of 10 and 5 nm were used, respectively.

#### 2.2.5 Guanidine hydrochloride and urea induced unfolding

Concentrations of GuHCl solutions were determined from the refractive index of the solution. As the concentration of the chemical denaturant increased, denaturation of the enzyme was observed at different pH conditions. To reach equilibrium, the peroxidase sample was incubated at the corresponding denaturant concentration for 24 hr at 25 °C. To obtain transition midpoints, as well as to judge the cooperativity or non-cooperativity of the transitions, the method used by Shirley (1995) was followed for the analysis of unfolding transitions. To compare the results from different measurements (molar ellipticity, fluorescence emission maximum, and relative activity), all the results were normalized and expressed as a fraction unfolded. The formula used for fraction unfolded calculation (Fu) is:

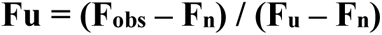

Where F_obs_ represent the observed signal at a given denaturant concentration, F_n_ represents the signal for the native protein, and Fu represents the respective signal of the denatured protein. The values for Fn and Fu are obtained by extrapolating the linear dependence of the signal on the concentration of the denaturant before and after the transition, respectively. Denaturant blanks are used to ensure that the observed signal is specific to the peroxidase and not due to the presence of the denaturant. This equation is used in a technique called spectroscopic denaturation, which is used to study the stability and unfolding of proteins.

#### 2.2.6 Thermal unfolding of peroxidase

Temperature-induced denaturation of the enzyme was performed under specified conditions as a function of increasing temperature. The enzyme samples are heated to increasing temperatures and kept at each temperature for 15 minutes before measurements are taken. The temperature of the samples is monitored using a thermocouple and a digital multimeter to ensure that the desired temperature is reached and maintained. Additionally, the samples are checked for protein aggregation using light scattering measurements, which can indicate if the proteins have undergone irreversible denaturation due to heat. This method allows for the study of how temperature affects the stability and folding of the enzyme, which can provide insight into its function and potential for thermal inactivation.

#### 2.2.7 Data analysis

The denaturation curves were plotted with the ratio of fluorescence intensity at the emission wavelength maxima of the native protein and the denatured protein against the denaturant molarity or temperature. A two-state N U unfolding mechanism is assumed, meaning that at any point on the curve, only the folded and unfolded conformations are present in significant concentrations. The method used by Shirley (Shirley, 1995) is followed to obtain the transition midpoints and judge the cooperativity or non-cooperativity of the transitions. For comparison of results obtained from different measurements such as molar ellipticity, fluorescence emission maximum, and relative activity, all the results are normalized and expressed as the fraction of unfolded protein. The fraction of unfolded protein, at any denaturant concentration, is calculated using the equation: fraction unfolded = (Fobs – Fn)/(Fu – Fn), as previously mentioned.

Denaturation curves of urea and guanidine hydrochloride (GuHCl) were monitored using CD spectroscopy. Denaturation experiments were conducted using Far-UV CD spectroscopy with an acquisition range of 215–260 nm. Data at 222 nm were used for fitting purposes, assuming a two-state model. This allowed for the determination of experimental equilibrium m (m_urea_ and m_GuHCl_) and C_m_ (concentration at which half-denaturation occurs) using following Equation:

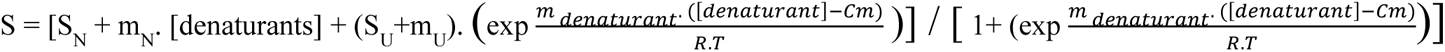

In this model, S represents the experimental CD signal as a function of denaturants concentration. S_N_ and S_U_ are the fitted CD signals for the Native and Unfolded states at 0 M denaturants, respectively. m_N_ and m_U_ represent the slopes of the native and unfolded state baselines, while m_denaturants_ describe the unfolding cooperativity. *R* is the ideal gas constant, and *T* is the experimental temperature (298.15 K) (Pacheco-García et al., 2022). This model provides a simplified approach for comparing the cooperativity of reversible chemical unfolding in *A. lakoocha* peroxidase.

## 3. Results

### 3.1 Absorbance spectroscopy

To elucidate the structure-function relationship of *A. lakoocha* peroxidase and its impact on enzymatic activity, various biophysical techniques were used. These approaches facilitate the observation of enzyme intermediates under varying pH conditions and during the denaturation process. The absorbance spectra of peroxidase showed four bands (Fig. 1A) in the visible and ultraviolet (UV) regions: the α-band at ∼630 nm, the β-band at ∼502 nm, the γ-band or Soret band at 404 nm, and the absorbance spectra of peroxidase at 280 nm. These bands can be used as a tool to investigate the structural and functional properties of *A. lakoocha* peroxidase. Changes in the wavelength or intensity of these bands upon ligand binding or unfolding can provide insight into the conformational changes of the enzyme. For example, substantial changes at 404 nm were observed upon ligand binding or unfolding, indicating changes in the tertiary and/or quaternary structure of the enzyme. However, changes at 280 nm were not observed under native or unfolded conditions, indicating that the secondary structure of the enzyme remains unchanged. The shifts at the 404 nm wavelength are evaluated to study the conformational change of the peroxidase. The absorbance spectra of *A. lakoocha* peroxidase in its native state, at pH 2 and in its unfolded state (6 M GuHCl) (Fig. 1B) are compared to observe the changes in the spectra. Under native conditions, the absorbance spectra show a soret peak at 404 nm, which is absent in the unfolded state. This indicates that the conformational change of the enzyme occurs upon unfolding. Additionally, Fig. 1C shows the peroxidase absorbance maxima measured at 404 nm under varying pH. The bell-shaped pattern suggests that the peroxidase is stable in the range of pH 3.5 to 8.5, which corresponds to the enzyme’s physiological pH range. This information is useful for understanding the stability and activity of the enzyme under different conditions.

**Fig.1.**
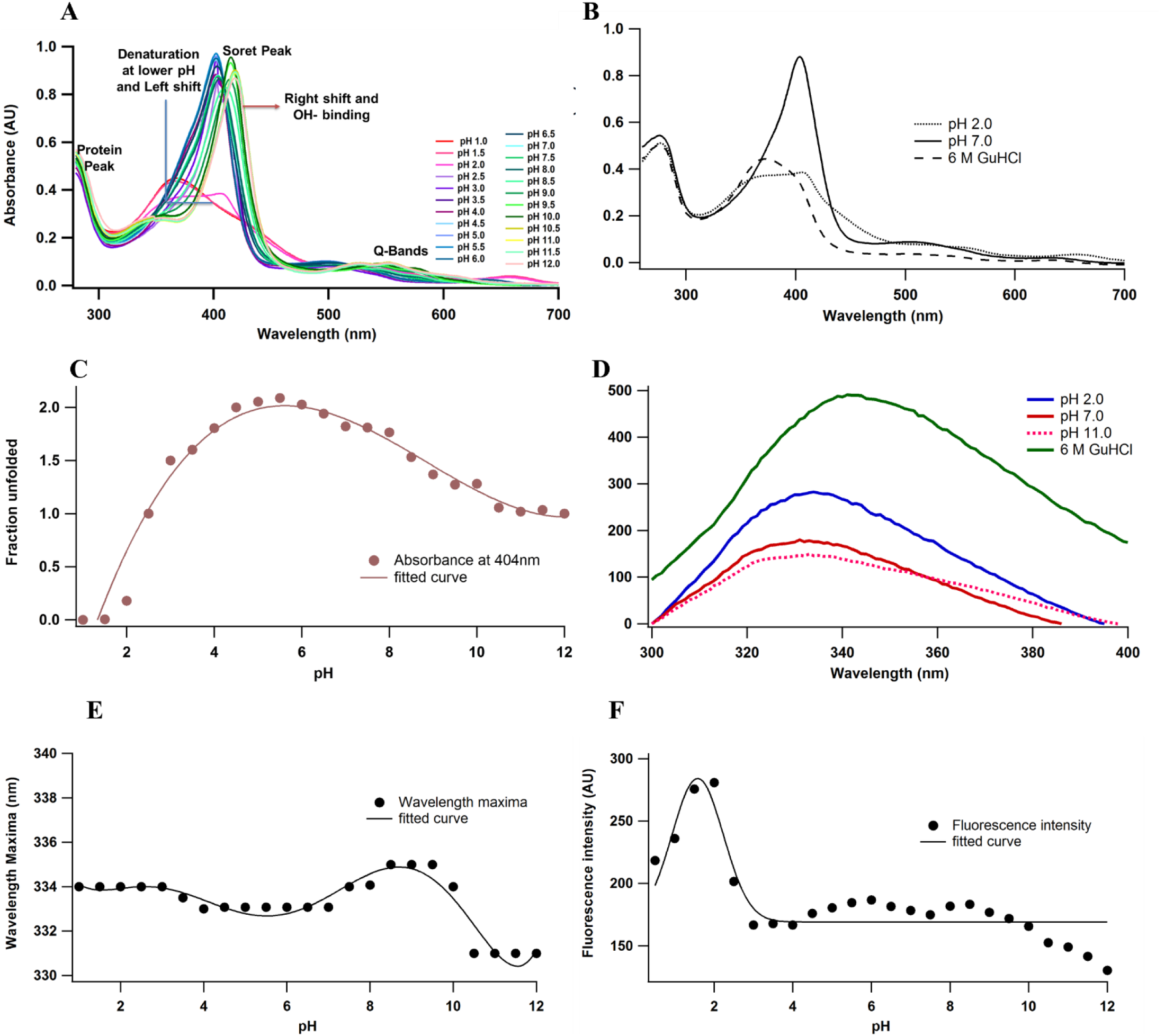
pH-induced conformational changes of *A. lakoocha* peroxidase: (A) UV absorbance spectra at different pH, (B) Spectra at pH 7.0 and at 6 M GuHCl have been compared with spectra at pH 2.0 (free of heme), (C) The absorbance spectrum at 404 nm under different pH conditions., (D) fluorescence spectra at different pH and in 6.0M GuHCl, (E) fluorescence wavelength maxima, and (F) fluorescence intensity at different pH.

### 3.2. Fluorescence spectroscopy

The fluorescence spectra of peroxidase under native conditions at pH 2, 11.0, and with 6 M GuHCl have been shown in Fig. 1D. The intrinsic fluorescence spectrum of peroxidase in its native state has an emission maximum at 333.07 nm, indicating a hydrophilic environment of its tryptophan residues. Under acidic and alkaline conditions, the fluorescence emission maximum remains the same, but compared to the native protein, the intensity of fluorescence increases by 57.4% at pH 2 (178.43 to 280.88) and decreases by 16.42% at pH 11 (178.43 to 149.13). This indicates that changes in pH can affect the microenvironment of the tryptophan residues and influence their fluorescence properties. On complete unfolding of the protein in the presence of 6 M GuHCl, the emission maximum of peroxidase suffers a red shift of 8 nm (333.07 nm to 341.07 nm) along with an approximately 175% increase in fluorescence intensity (from 178.43 to 491.04) compared to the native condition. This suggests that upon unfolding, the tryptophan residues are exposed to a more hydrophobic environment, resulting in an increase in fluorescence intensity and a red shift in the emission maximum. The effect of different pH on *A. lakoocha* peroxidase was also observed by the intrinsic fluorescence emission scale. The wavelength maximum of peroxidase as a function of pH has been shown in (Fig. 1E). The transition curve did not show much change in the emission maximum under varying pH and appeared to occur through three state transitions. The first transition shows a red shift of 1 nm on decreasing pH from 4.0 to 2.0 under acid unfolded state. The second transition occurred between 6.0 and 8.5 pH along with a red shift of 2nm. On increasing the pH to the basic region, a third transition occurs in the range of 9.0 to 10.5 pH, with a blue shift of 4 nm in the fluorescence emission maxima, resulting in the alkaline denatured state of peroxidase. The pH-induced transition is non-cooperative with a slight change in the transition maxima because tryptophan residues, even in their native state, seem to be exposed to the solvent. Changes in the fluorescence intensity as a function of pH are shown in Fig. 1F. The fluorescence intensity spectra displayed two transition phases. The first transition occurs from the native state to acid unfolded state (between pH 4.0 to pH 2), with a transition mid-point at pH 3.0, where the fluorescence intensity is increased marginally. The second transition towards basic pH is incomplete and takes place from pH 8.0 to 12.0. This study of the intrinsic fluorescence spectra of peroxidase under different pH conditions provides insight into the conformational changes occurring in the enzyme and its stability at different pH.

### 3.3. ANS binding assay

ANS (8-anilino-1-naphthalene sulphonate) is also known as a polarity-sensitive extrinsic fluorescent probe. The effect of pH on the ANS fluorescence of peroxidase at different pH is shown in Fig. 2A and B. ANS emission spectra were recorded with excitation at 380 nm. At pH 2.5 proteins show partial ANS binding. On further decreasing the pH from 2.5 to 1.0, an increase in fluorescence intensity was observed. At pH 2 enzymes showed maximum ANS binding. Compared to native state, at pH 2 ANS fluorescence intensity showed up to 5-fold increase and fluorescence maxima exhibited the red shift from 481 nm (pH 2) to 508.05 nm (pH 7). ANS fluorescence decreases on either side of 2 pH and at the alkaline denatured state (UB state, pH 8-12), ANS binding is almost absent and stable up to pH 12.

**Fig.2.**
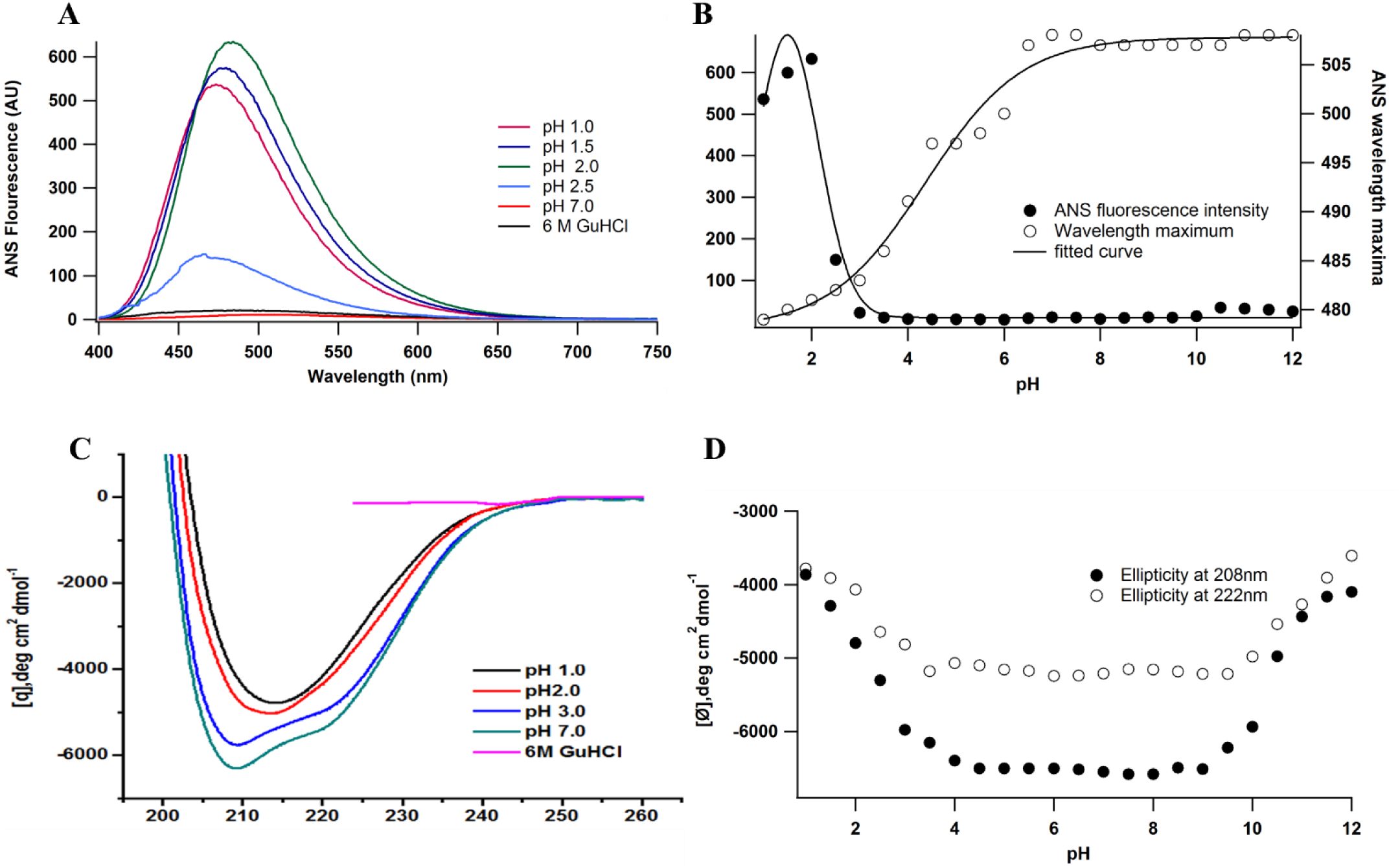
Effect of pH on the ANS fluorescence of *A. lakoocha* peroxidase: (A) ANS binding to peroxidase was studied at pH 1.0, pH 1.5, pH 2.0, pH 2.5, pH 7.0, and in the presence of 6M GuHCl, and (B) For intrinsic fluorescence measurement, samples were incubated for 24 h at 25oC. ANS emission spectra were recorded with excitation at 380nm., pH induced conformational changes of *A. lakoocha* peroxidase: (C) far-UV CD spectra at different pH and in 6.0 M GuHCl, and (D) Ellipticity at 208 nm and 222 nm as a function of pH.

### 3.4. Circular dichroism

The effect of varying pH on the secondary structure of peroxidase was also studied. The CD spectra of *A. lakoocha* peroxidase at acidic, neutral, and denatured conditions (6 M GuHCl) are presented in Fig. 4A. At pH 7 in the far-UV region, the CD spectrum of peroxidase comprises strong negative ellipticity at 208 and 222 nm, suggesting that the molecule may consist of α-helix and β-sheet rich regions and belong to the α + β class of proteins (Kumar, Tripathi, de Moraes, Caruso, & Jagannadham, 2014; Manavalan, 1983). Fig. 2C displays that the mean residue ellipticity at 222 nm is −5300.2 deg.cm^2^ dmol^-1^ in the native condition (Fig. 4A) while under an acidic state at 1.0 pH, the ellipticity at 222 nm shifts to −3779.9 deg.cm^2^ dmol^-1^. However, under similar conditions, all the spectral features of peroxidase are lost in the presence of 6 M GuHCl, indicating that the enzyme drops the secondary structural organization under denaturing conditions.

The structural changes of peroxidase according to pH are indicated by shifts in ellipticity at 208 and 222 nm, as shown in Fig. 2D. These structural changes display that the secondary structural contents of proteins remain the same in the range of 4.0-9.5 pH and decrease on either side of this range, without changing the shape of the spectra. The changes are similar in magnitude at lower pH and in the basic range.

### 3.5. Guanidine hydrochloride induced unfolding of peroxidase

Guanidine hydrochloride (GuHCl) induced the unfolding of peroxidase under neutral conditions, which, when examined by alterations in secondary structure, absorbance, and fluorescence, showed structural stability till 3.0 M GuHCl (Fig. 3A). Under natural conditions, GuHCl-induced unfolding results in cooperative sigmoidal curves (Fig. 5A) with a transition midpoint at 4.5 ± 0.1 M GuHCl. However, at pH 2, enzymes displayed structural stability up to 1.5 M GuHCl (Fig. 3B). The transitions as measured by different spectroscopic techniques are cooperative and non-coincidental (Fig. 3B). This observation illustrated that fluorescence intensity is lost first, followed by losses in secondary structure. Loss in secondary structure took place between 2.0 and 5.0 M of GuHCl, whereas the changes in intrinsic fluorescence occurred between 1.5 and 5.0 M of GuHCl at pH 2. The transition mid-points (Cm) of denaturation followed by far UV CD are 3.5 ± 0.1 M, and by absorbance and fluorescence intensity is approximately 3.25 ± 0.1 M. The effect of GuHCl on the alkaline denatured state of a protein at pH 11 is similarly studied. The transitions as measured by different spectroscopic techniques are cooperative and coincidental (Fig. 3C). Loss of secondary structure and intrinsic fluorescence took place between 2.5 and 5.0 M of GuHCl, and the enzyme showed structural stability until 2.0 M of GuHCl.

**Fig.3.**
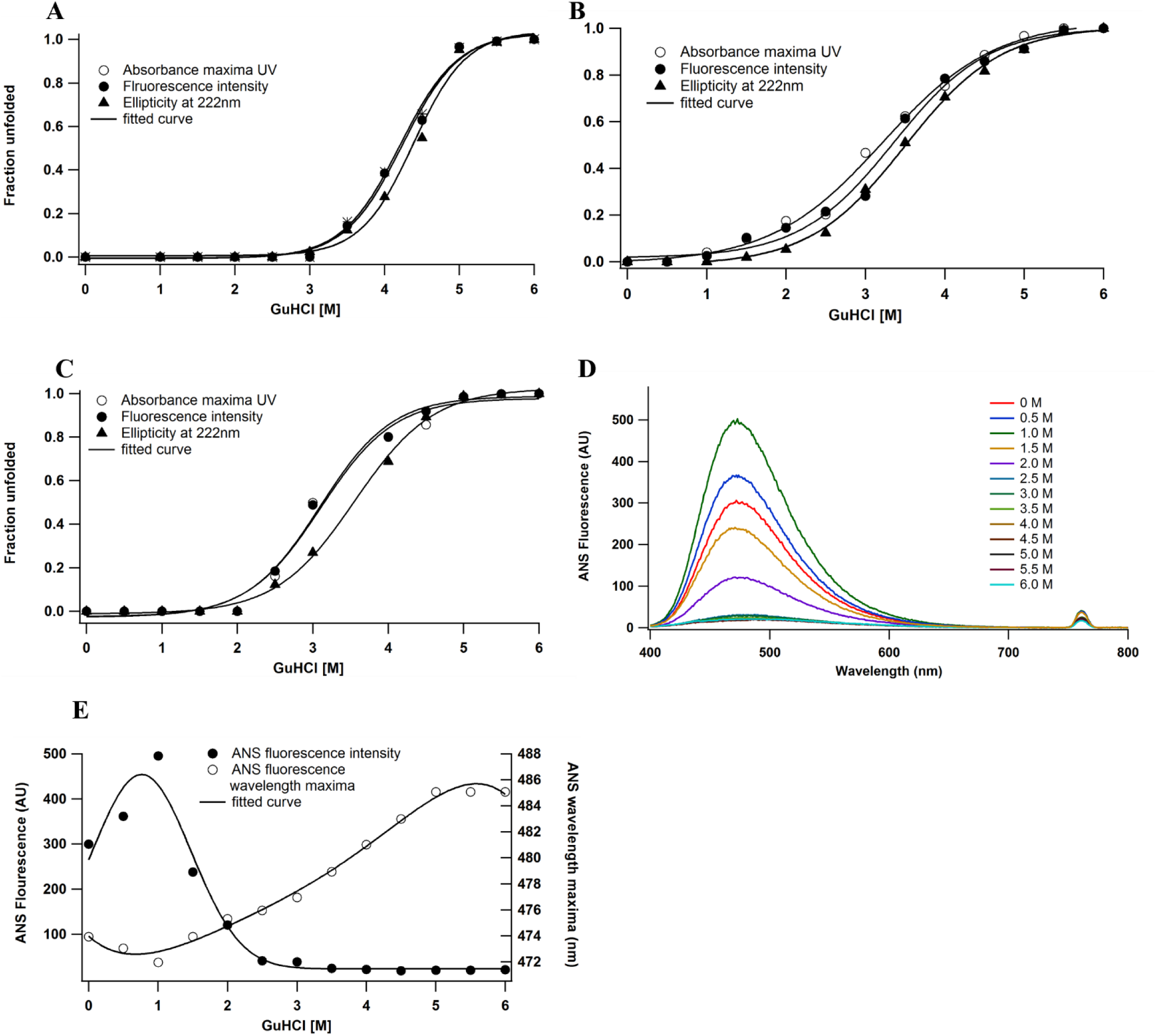
GuHCl induced unfolding of A. lakoocha peroxidase at different pH: (A) pH 7.0, (B) pH 2.0, and (C) pH 11.0, by following far-UV CD, fluorescence, and UV absorbance, Effect of pH on the ANS fluorescence of A. lakoocha peroxidase: (D) ANS binding spectra, as a function of increasing concentration of GuHCl at pH 2.0 (E) ANS fluorescence intensity and wavelength maxima at pH 2.0, as a function of increasing concentration of GuHCl.

When ANS fluorescence was used to monitor the denaturation process in the native and alkaline denatured states at pH 11, peroxidase did not specify any ANS binding, even after a prolonged exposure. That may be due to an absence of contact with the hydrophobic surface. These characteristics indicate that *A. lakoocha* peroxidase is more susceptible to denaturation in acidic conditions in comparison to alkaline conditions. So, it is meaningful to observe the structural integrity of the molecule at an acidic pH. Fig. 3D shows the ANS binding pattern of peroxidase with increasing concentrations of GuHCl, at pH 2. Under the acid denatured state, a decrease in ANS fluorescence intensity by increasing the molarity of GuHCl was observed with a red shift in its wavelength maximum with GuHCl at pH 2 (Fig. 3E).

### 3.6. Urea induced unfolding of peroxidase

Under neutral conditions, urea does not cause any structural agitation in the protein, as the enzyme retains its maximum structural and functional parameters even at 5.5 M urea concentration (Fig. 4A). The urea induced unfolding studies are therefore carried out at low and high pH, where enzymes are more susceptible to denaturants. However, urea induced unfolding at a lower pH yielded some interesting facts. The denaturation is followed by absorbance, far UV CD and fluorescence. At pH 2, the urea induced unfolding is cooperative, but non-coincidental (Fig. 4B). The transition mid-points for intrinsic fluorescence and absorbance are 4.25 ± 0.1 M and for secondary structure are 4.5 ± 0.3 M. The loss in secondary structure occurred between 2.0 and 7.0 M urea. But, at pH 11, where the enzyme is in an alkaline denatured state, the unfolding transitions by different probes are cooperative, with a transition mid-point at 4.75 ± 0.1M (Fig. 4C).

**Fig.4.**
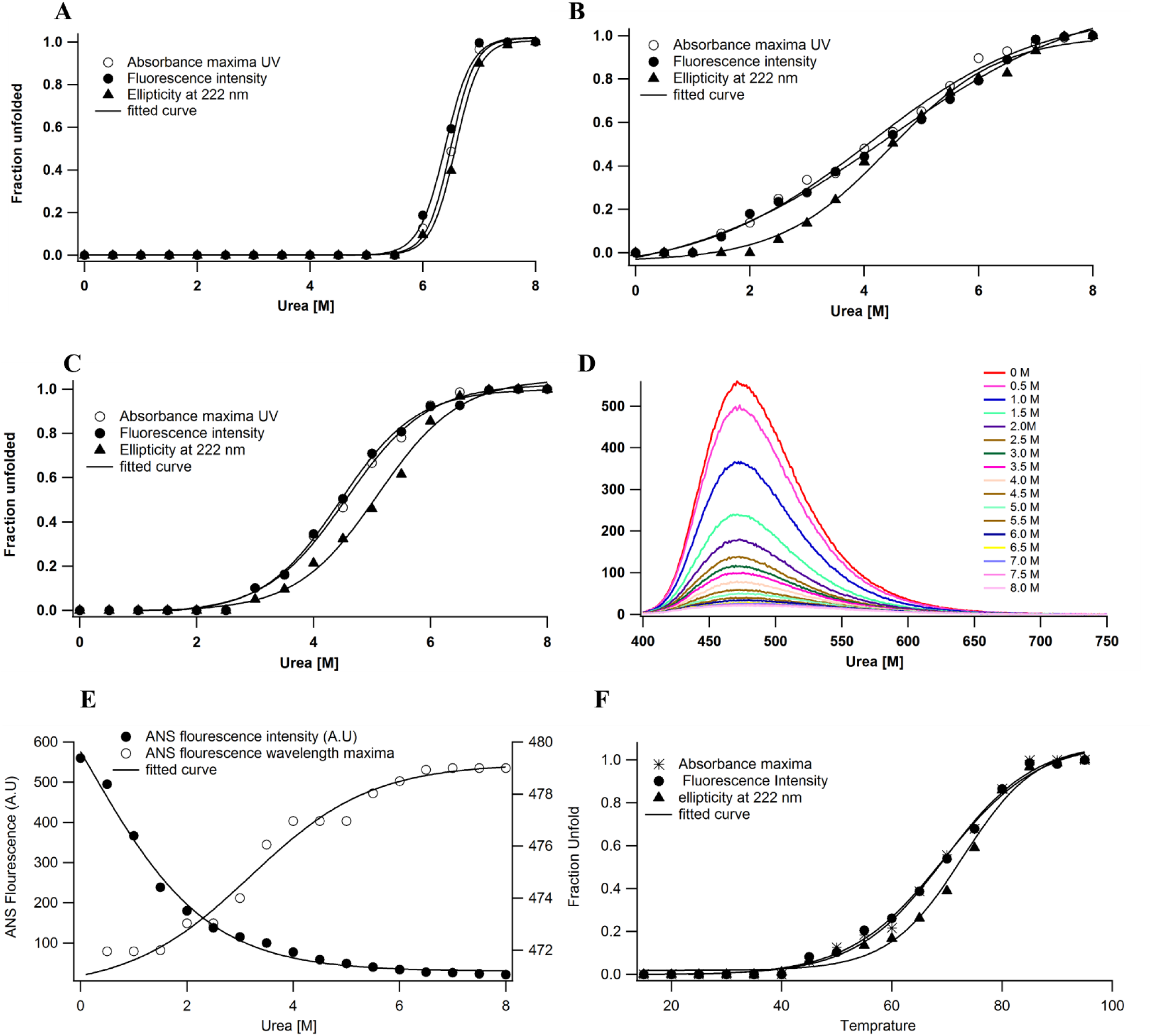
Urea induced unfolding of A. lakoocha peroxidase at different pH: (A) pH 7.0, (B) pH 2.0, and (C) pH 11.0, by following far-UV CD, Fluorescence and UV absorbance, Effect of pH on the ANS fluorescence of A. lakoocha peroxidase: (D) ANS binding spectra, as a function of increasing concentration of urea at pH 2.0, (E) ANS fluorescence intensity and wavelength maxima at pH 2.0, as a function of increasing concentration of urea, and (F)Temperature induced unfolding of A. lakoocha peroxidase at pH 7.0: The structural changes are followed by UV absorbance, far-UV CD, and fluorescence.

The cooperative but non-coincidental transitions at pH 2, along with an increase in ANS binding at low concentrations of urea (Fig. 4D and E) indicate that a discrete intermediate state exists in the urea induced unfolding of peroxidase at pH 2. An increase in ANS fluorescence intensity coupled with a red shift in the wavelength maximum of peroxidase at pH 2 is shown in Fig. 4D and E.

### 3.7. Thermal denaturation of peroxidase

Temperature, the oldest mode of protein denaturation, can induce structural changes and hence provide ample information about a protein molecule. The temperature induced unfolding of peroxidase at pH 7 was measured (Fig. 4F). The transition midpoints of temperature induced unfolding of peroxidase are very high and come in the range of 62.5 ± 0.5 °C. Further, the structure of the protein melts at a much higher temperature, at 70 °C, indicating marked structural agitations already induced by the increased concentration of protons.

## 4. Discussion

In the era of proteomics, it is becoming very important to understand the structure-function relationship of proteins using various biophysical tools (Neet & Lee, 2002). Despite the importance of peroxidase, relatively few biophysical studies have been carried out on this class of proteins. Because of the dependency of catalytic activity on pH, the stability of an enzyme determines its functionality and may limit its wide application. Therefore, pH denaturation of peroxidase was monitored by various spectroscopic techniques to probe the different aspects of conformational changes in its structure and the identification of populated intermediates at extremes of pH, during denaturation.

### 4.1. Effect of pH on the absorbance spectra of A. lakoocha peroxidase

The absorbance technique is used to measure the absorption of radiation by a sample as a function of wavelength or frequency. The absorbance spectra of the protein at different pH levels are shown in Fig. 1A, which shows spectra of the protein under neutral conditions that are similar to the characteristic heme peroxidase spectra. The absorption bands in the visual spectrum region (α- and β-peaks) and at the boundary of the visual and UV parts (Soret γ) are due to the intensively colored prosthetic groups of peroxidases. Other absorption bands of peroxidase solution are located in the UV region: φ-band with a maximum of 280 nm, indicating a significant contribution from tryptophan and tyrosine residues also due to the light absorption by aromatic amino acids in proteins. As revealed from Fig. 1A, peroxidase spectra do not show much change in the range of pH 2.5 to 8.5, which indicates good stability of peroxidase in this state. But at lower pH, heme peroxidase spectra express a left shift with a declined absorbance at 404 nm due to denaturation, which is a clear indication of free heme in the range of pH 1 to 2. This result specifies that at lower pH peroxidase releases its prosthetic group (heme). Whereas beyond pH 8, the heme group of peroxidase shows OH^-^ or H_2_O binding, with an increase in pH, which results in the right shifting of soret band. At higher pH, the Q-band of peroxidase gets split and the soret peak shifts to the right. Though the shift in absorbance intensity at the 404 nm (Soret peak) gives information regarding the stability and activity of peroxidase, this shift at 404 nm wavelength is investigated further to follow the conformational change of the protein in the present investigation.

The spectra in Fig. 1B show a decrease in the soret band of peroxidase at pH 2 and 6 M GuHCl, indicating the loss of the heme group and denaturation of the protein. Fig. 1C shows that the enzyme retains proteolytic activity in the pH range of 3.5-8.5 and drops on either side of it, suggesting that the native structure and functional stability of the protein are maintained in this pH range. The decline in proteolytic activity outside of this range is likely due to the repulsion of columbic forces from the positive charges of the polypeptide chains.

### 4.2. Fluorescence spectroscopy

The fluorescence emission maximum of chromophores, such as tyrosine and tryptophan residues, can change when exposed to different solvent environments. The quantum yield of fluorescence can also decrease when the chromophores interact with quenching agents in the solvent or within the protein. In the case of *A. lakoocha* peroxidase, the fluorescence spectra under different pH and denaturing conditions are shown in Fig. 1D. The tryptophan excitation spectra reflect the average environment of the tryptophan and shows a red shift in the emission maximum when the chromophores are exposed to solvent. The results of the experiment, as observed in Fig. 1E, indicate that the tryptophan residues become exposed to solvents due to the unfolding and disclosure of chromophores to the solvent, particularly under denaturing conditions with 6 M GuHCl. Additionally, the intensity of fluorescence increases at pH 2 but decreases at pH 11, likely due to the presence of a heme group in the peroxidase that acts as a main quencher and quenches the energy in the native state. However, upon denaturation with 6 M GuHCl, the fluorescence intensity is amplified compared to the acidic condition, indicating the complete exposure of tryptophan residues to an aqueous environment.

The observations suggest that the fluorescence intensity of the peroxidase protein increases as the protein loses iron upon denaturation. The pH-induced transition of peroxidase in the alkaline range, as monitored by fluorescence intensity, seems to be cooperative, with a transition mid-point at pH 3. The fluorescence wavelength maxima remained largely unaffected throughout different pH values (Fig. 1E), but the fluorescence intensity revealed changes at different pH values (Fig. 1F). Lower pH values resulted in higher fluorescence intensity, while higher pH values resulted in a decrease in fluorescence intensity, likely due to the quenching of tryptophan fluorescence by specific quenching groups such as protonated carboxyl and imidazole groups.

### 4.3. Acid-induced changes in peroxidase

ANS binding is a tool used to measure the exposure of hydrophobic surfaces in enzymes. The magnitude of ANS binding can indicate the structural integrity of a protein. In the case of *A. lakoocha* peroxidase, at pH levels above 3, no binding of ANS to peroxidase was observed. However, at lower pH levels, some binding was detected, albeit with a lower magnitude. As the pH decreased from 3 to 2.5, an increase in ANS binding was observed, as indicated by an increase in fluorescence intensity from 21.76 to 149.78 and a blue shift of the emission maximum from 483 nm to 482 nm. On further decreasing pH to lower values, the protein gets refolded by pH 0.5, driving the molecule into acid refolded state caused by the shielding of columbic repulsions by counterions., as shown in Fig. 2A. However, at pH 2 proteins show maximum ANS binding along with a significant amount of secondary structure with no tertiary contacts. This sudden increase in ANS fluorescence intensity is due to maximum exposure of hydrophobic surfaces. That is one of the properties of molten globule state. This result indicates the presence of molten globule state at pH 2. The results of the experiment as observed in Fig. 2A show that the peroxidase retains its proteolytic activity and native structure within a pH range of 3.0-10.0, as indicated by the absence of ANS binding in this range, suggesting that the protein has a relatively stable and rigid native conformation in a wide range of pH values but becomes more flexible and susceptible to changes in acidity.

The pH-induced conversion of this protein shows the first transition phase in between and pH 2.5 and 4.0 (midpoint 3.25 plus second transition in the middle of pH 1 and 2.5 (midpoint 1.5). As observed by absorbance and fluorescence spectra, at pH 7.5 and above, protein turns into an alkaline denatured state (UB), besides at pH 1.5 or lower, the protein unfolds to form acid-unfolded state (UA). While on further decreasing the pH to 0.5 induces refolding of the UA-state and the resulting state has been termed as acid-refolded ‘A-state’ (Fig.2).

#### Circular dichroism

The use of circular dichroism (CD) spectroscopy and the hydrophobic dye ANS allowed for the study of the structure and stability of *A. lakoocha* peroxidase. The Far-UV CD spectra of the enzyme (Fig. 2C) showed it to be consistent with the α + β class of proteins and possibly contain distinct α and β-rich domains. The peroxidase was found to be quite stable in terms of secondary structure, with only a 50% decrease in ellipticity at lower pH. However, under denaturing conditions, the peroxidase lost all prominent peaks in aromatic regions, indicating the loss of secondary structural features and the enzyme being in an unfolded state. The shifts in ellipticity at varying pH were found to not follow a simple two-state transition but rather a three-step transition (Fig. 2D). The first transition from the native state to acid unfolded state occurred between pH 4 - 1, with a half bell shape in the range of pH 1 - 4, revealing the loss of some prominent peaks. This loss of secondary structure, as well as reduced activity, represents the enzyme acid unfolded state. The second transition occurred between pH 4 to 8 under native conditions, and the third transition occurred between pH 8 – 12 with a mid-point at 10.5 pH. The use of hydrophobic dye ANS also revealed that despite the melting of its secondary structure, a large amount of hydrophobic clusters were left over in the acid-unfolded state (Fig. 2A).

### 4.4. GuHCl induced unfolding of peroxidase

The analysis of denaturation curves induced by GuHCl can provide information about the conformational stability of the enzyme. The denaturation of peroxidase by GuHCl at neutral pH is a cooperative, two-state process with a high transition mid-point of about 4.5 ± 0.1M (Fig. 3A). However, at pH 2 and 11.0, (Fig. 3B and C) the transition mid-points are lower as compared to the same parameters at pH 7. When ANS fluorescence was used to monitor the denaturation process under different conditions, enzyme only reveals ANS binding with acidic condition. Where the decrease in ANS fluorescence intensity coupled with red shift in its wavelength maximum was found, as shown in fig.3D. At pH 2 maximum ANS binding as well as other spectroscopic properties exist due to increase of distance from the heme group, which serves as a quencher in heme containing peroxidase. This kind of quenching effect of heme has also been reported in other heme containing peroxidases, like lactose peroxidase and horse radish peroxidase (Carvalho et al., 2003; Zelent B, 2010). ANS binding to peroxidase at pH 2 in the presence of ≥2.5M GuHCl suggests the absence of surface hydrophobic patches, since no ANS binding was observed. While decreasing the concentration of GuHCl, one intermediate was found at 1.0 M GuHCl, here the protein shows maximum (Fig.3E) ANS fluorescence intensity, which indicates the exposure of hydrophobic residues in this state. That’s in line with the idea of molten globule state (MG) at the particular GuHCl concentration. While there is absence of ANS binding to peroxidase at 3 M GuHCl and above, which suggests the absence of surface hydrophobic patches and refolding and stabilization due to specific stabilizing interactions between the intermediate state and GuHCl.

### 4.5. Urea induced unfolding of peroxidase

The stability of the enzyme peroxidase against urea was evaluated under different pH conditions. Results revealed that at neutral pH, the enzyme retained its native structure and function even when exposed to 5.5M urea (Fig.4A), indicating stability against urea. However, at an acidic pH (2), urea caused the enzyme to unfold cooperatively, with transition midpoints for secondary and tertiary structures at 4.5 ± 0.3M and 4.25 ± 0.1M urea, respectively (Fig. 4B). The unfolding process was observed using far-UV CD and fluorescence, which revealed cooperative and non-overlapping transitions (Fig. 4B) along with an increase in ANS binding at low concentrations of urea (Fig. 4D & E). Additionally, ANS fluorescence intensity, which is highest at pH 2 in the absence of denaturant, decreased parallel with an increase in urea concentration. The sigmoidal change in fluorescence intensity with urea at pH 2 is attributed to configurational changes in the amino acids. The unfolding process at pH 2 showed the same changes in far-UV CD and wavelength maxima as at pH 7 (Fig. 4B). However, complete loss of activity above 5 M urea, a sigmoidal increase in fluorescence intensity, ANS binding, and conversion to beta-sheet-like spectra seen by far-UV CD above 3.5 M urea were due to perturbation of the enzyme’s functional structure (Fig. 4B). These results suggest that a discrete intermediate state exists in the urea-induced unfolding of peroxidase.

Overall, this study found that the enzyme peroxidase unfolds differently when exposed to urea and guanidine hydrochloride (GuHCl) under similar conditions. The results showed that urea is less effective than GuHCl in reducing the mid-points of unfolding at a given concentration. This difference in behavior is likely due to the ionic character of the two denaturants. Urea and GuHCl interact with peptide bonds but have different electrostatic and hydrophobic properties. GuHCl, which is positively charged, may stabilize the protein through its anionic effect, while urea does not have this effect. Additionally, the counter-anion of GuHCl, chloride, may shield electrostatic repulsion and allow forces that favor the folding and stability of the protein. However, the stabilizing effect of GuHCl may not always be predominantly ionic in nature (Zarrine-Afsar, Mittermaier, Kay, & Davidson, 2006). The study also discussed the effects of temperature on the structure and activity of peroxidase from *A. lakoocha*. It was found that the enzyme retained approximately 45% of its structure under neutral conditions and had a high degree of stability, with a transition midpoint (Tm) of 62.5±0.5°C (Fig. 4F). The complete unfolding parameters of *A. lakoocha* peroxidase are summarized in Table 1. The accumulation of intermediates in the GuHCl and urea induced unfolding pathways is presented here, which indicates that peroxidase may possess two domains that are behaving differently in various denaturing conditions, leading to the formation of a number of intermediate states.

**Table.1:** Unfolding parameters of *A. lakoocha* peroxidase.

| Condition | Denaturant | Method | Transition mid-point |
| --- | --- | --- | --- |
| pH 7.0<br>(Native state) | GuHCl | CD[$\Theta$]<sub>222</sub> | $4.5 \pm 0.1$ M |
| | | Fluorescence | $4.5 \pm 0.1$ M |
| | | absorbance | $4.5 \pm 0.1$ M |
| | Urea | CD[$\Theta$]<sub>222</sub> | $6.5 \pm 0.1$ M |
| | | Fluorescence | $6.5 \pm 0.1$ M |
| | | absorbance | $6.5 \pm 0.1$ M |
| | Temperature | CD[$\Theta$]<sub>222</sub> | $65.0 \pm 0.5^\circ\text{C}$ |
| | | Fluorescence | $62.5 \pm 0.5^\circ\text{C}$ |
| | | absorbance | $62.5 \pm 0.5^\circ\text{C}$ |
| pH 2.0<br>(Molten globule state) | GuHCl | CD[$\Theta$]<sub>222</sub> | $3.5 \pm 0.1$ M |
| | | Fluorescence | $3.25 \pm 0.1$ M |
| | | absorbance | $3.25 \pm 0.1$ M |
| | Urea | CD[$\Theta$]<sub>222</sub> | $4.5 \pm 0.3$ M |
| | | Fluorescence | $4.25 \pm 0.1$ M |
| | | absorbance | $4.25 \pm 0.1$ M |
| pH 11.0<br>(Alkaline denatured state) | GuHCl | CD[$\Theta$]<sub>222</sub> | $3.5 \pm 0.1$ M |
| | | Fluorescence | $3.5 \pm 0.1$ M |
| | | absorbance | $3.5 \pm 0.1$ M |
| | Urea | CD[$\Theta$]<sub>222</sub> | $4.75 \pm 0.1$ M |
| | | Fluorescence | $4.75 \pm 0.1$ M |
| | | absorbance | $4.75 \pm 0.1$ M |

## 5. Conclusion

In this study, we observed three different folding states of a peroxidase from the latex of *A. lakoocha* using various spectroscopic techniques under different pH, denaturing conditions, and temperatures. At pH 7, the enzyme is in its **native state,** with proteolytic activity, intact secondary structure, and without any hydrophobic patches. At lower pH values, the enzyme transitions through intermediate states such as the **molten globule** state, the acid-unfolded state (**UA state**) and acid refolded state **(A state)**. In the alkaline denatured state **(UB state**), ANS binding is almost absent, indicating that *A. lakoocha* peroxidase is more susceptible to denaturation under acidic conditions, than alkaline conditions. Fluorescence intensity decreases at higher pH in the UB state due to a reduced distance between tryptophan and specific quenching groups, leading to quenching of tryptophan fluorescence. Under denaturing conditions, the enzyme can reach a completely denatured state when exposed to GuHCl concentrations above 5 M. At low concentrations (1.0 M), GuHCl acts as an anion, while at concentrations of 2.5 M or higher, it behaves as a chaotrope. Further experiments revealed that incubation of the peroxidase in 8 M urea overnight resulted in only a 60% loss of secondary structure, whereas incubation with 6 M GuHCl led to a complete loss of secondary structure. Our study demonstrates that the purified peroxidase from the latex of *A. lakoocha* is a robust model system for investigating protein folding pathways. The identification of these intermediate states represents a significant advancement in the field of protein folding and lays the foundation for future research into the enzyme’s physiological substrates, structural characteristics, and potential applications in the biotechnology and pharmaceutical industries.

## Author Contribution

K.S. purified the protein and performed biophysical experiments, with assistance from M.P. and A.K. for data analysis. The experimental design was planned by K.S. and J.M. The manuscript was written by K.S. with input from all authors. We thank Prof. Suman Kundu from the Department of Biochemistry at UDSC Delhi for providing the necessary infrastructure.

## Conflict of interest

The authors declare no competing financial interests.

## Acknowledgements

Financial assistance from the CSIR, Government of India, in the form of fellowships to KS is acknowledged. Grants and funds from UGC and DBT for infrastructure are also acknowledged.

## Notes

### Competing Interest Statement

The authors have declared no competing interest.

### Summary of Updates

Figures are revised.

